# *Toxoplasma gondii* in house mice *Mus musculus domesticus* from Dakar, Senegal: clues for oral rather than vertical mode of infection

**DOI:** 10.1101/605014

**Authors:** Lokman Galal, Claire Stragier, Farid Boumediène, Azra Hamidović, Océane Maugrion, Marie-Laure Dardé, Carine Brouat, Aurélien Mercier

## Abstract

*Toxoplasma gondii* is an ubiquitous highly prevalent zoonotic protozoan. Cats are the definitive hosts, while all other warm-blooded animals are intermediate hosts for this parasite. Commensal rodents, being the main preys of cats, are probably the major reservoir for *T. gondii.* Rodents often develop dormant tissue cysts after ingestion of oocysts shed in the environment by cats in the form of contaminated feces. Experimental evidence that vertical transmission can be sufficient to the perpetuation of transmission between generations of mice has also been found. In natural settings, the relative epidemiological importance of vertical transmission over oral transmission is a matter of debate and raises the question of the possibility of a *T. gondii* cycle in the absence of cats. In the present study, we took advantage of an extensive survey of commensal rodents in Dakar, Senegal, where the house mouse *Mus musculus domesticus* is the predominant putative reservoir of *T. gondii.* Host genotype and spatial location through GPS referencing of all trapping localizations were investigated in relation to *T. gondii* infection in 12 sites of the city of Dakar and on Goree Island. In each sampled site, the occurrence of over-prevalence zones of *T. gondii* infection was investigated through Kulldorf’s statistic using SaTScan software. For the sites where a possible over-prevalence zone was identified, mice family lines were inferred from Discriminant Analysis of Principal Components (DAPC). In 3 of the 4 identified possible over-prevalence zones, *T. gondii* infection was not confined to a single family line, which suggested no association between kinship and infection occurrence. This finding rather suggests an environmental source of infection for mice associated with localized putative foci of environmental contamination and supports an oral route of infection for mice from Dakar rather than a cycle based on vertical transmission.

## 1. Introduction

Toxoplasmosis is an ubiquitous highly prevalent parasitic zoonosis, caused by an obligate intracellular protozoan parasite *Toxoplasma gondii.* In human, *T. gondii* infection is often subclinical, except in some risk groups like developing fetus (in case of congenital infection) and immunocompromised patients, for which toxoplasmosis can have severe health consequences. Felids are the definitive hosts, with the domestic cat being the unique definitive host in the domestic environment, while all other warm-blooded animals are intermediate hosts for this parasite. Birds and mammals, including humans, often develop dormant tissue cysts after ingestion of oocysts shed in the environment by cats in the form of contaminated feces. Commensal rodents, being the main preys of cats (Turner and Bateson, 2013), are probably the most important reservoirs for *T. gondii* in the domestic environment.

Experimental studies showed that rodents are able to get infected by *T. gondii* through several routes (Dubey, 2009). Infection through oocyst ingestion is universally admitted as a conventional infection route for most of intermediate host species including rodents, although a high inoculum dose can be fatal in some susceptible hosts (Owen and Trees, 1998). The occurrence of vertical transmission appears to vary between species of rodents, but also between the different lineages within a single species (Hide, 2016). Beverley (1959) showed that vertical transmission can maintain *T. gondii* infection within a single line of outbred laboratory mouse over 9 generations without exogenous re-infection. In contrast, in BALB / c mice, hamsters, and in most of the laboratory rat lineages, vertical transmission could be noticed mainly when the infection occurs during pregnancy, but seldom in individuals with a chronic infection (Dubey et al., 1997; Freyre et al., 2009; Roberts and Alexander,1992).

An important role of vertical transmission in intermediate hosts, bypassing sexual multiplication in the definitive host, was proposed as a possible explanation for the clonal structure of most *T. gondii* populations (Worth et al., 2013). However, the ability of *T. gondii* to modify the behavior of rodents to presumably facilitate their predation and their trophic transmission to cats (Webster et al., 1994; Berdoy et al., 2000; Gonzalez et al., 2007; Vyas et al., 2007) suggests that oral transmission is of key importance in this current pattern of host-parasite coevolution. In natural settings, the relative epidemiological importance of vertical transmission over oral transmission is challenging to test and is a matter of debate (Dubey, 2009). Estimating the epidemiological importance of vertical transmission based on the levels of prevalence in embryos collected from pregnant dams could face some limits. Indeed, Beverley (1959) noticed that experimental congenital infection in mice is associated with heavy mortalities among the offspring and does significantly reduce the number reared to maturity in each litter.

In this study, the transmission pathways of *T. gondii* were indirectly investigated in mice *(Mus musculus domesticus)* from Dakar, the capital of Senegal (West Africa) and the biggest city of this country. Our objective was to examine whether *T. gondii* infection occurrence was explained by the degree of kinship between mice, in order to evaluate the likelihood of vertical infection in the wild.

## 2. Materials and methods

We took advantage of an extensive survey of urban rodents in the city of Dakar. Briefly, sampling was carried out in 12 sites of the Cape Verde Peninsula and in the Gorée Island (Figure 1), each site being separated from each other by a minimum distance of 600 m and covering a median surface of 0.04 km^2^. Two traps (one wire mesh trap and one Sherman trap) were set per room or courtyard in buildings corresponding to dwelling houses, boutiques, workshops, offices or warehouses, and whose locations were precisely recorded with a GPS device. This survey led to the sampling of 481 mice, which were genotyped using a set of 15 microsatellite markers (Stragier et al., 2019) and whose the infectious status was determined using a real-time PCR specific to *T. gondii* on brain DNA extracts (Galal et al., 2018b).

**Figure 1.**
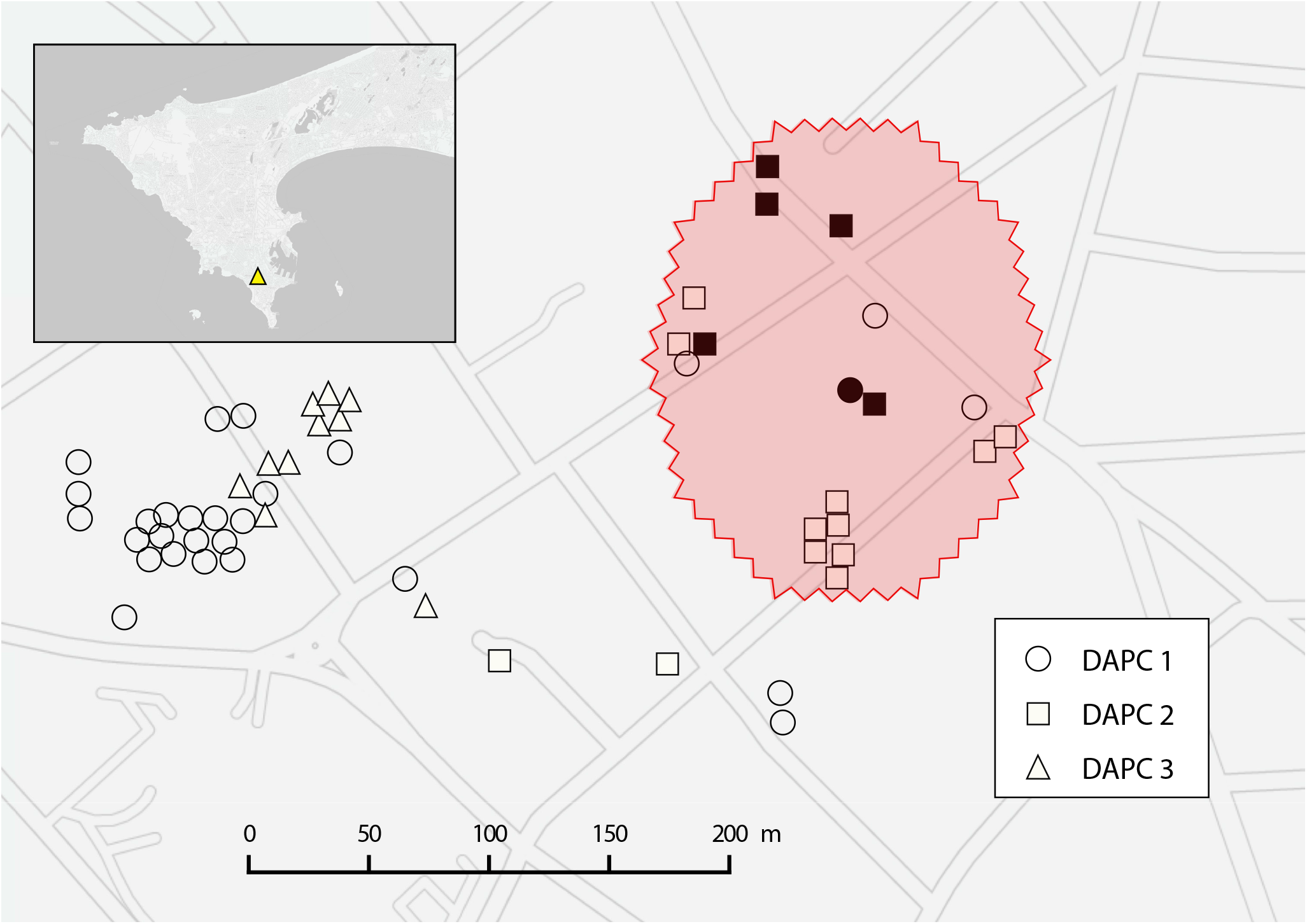
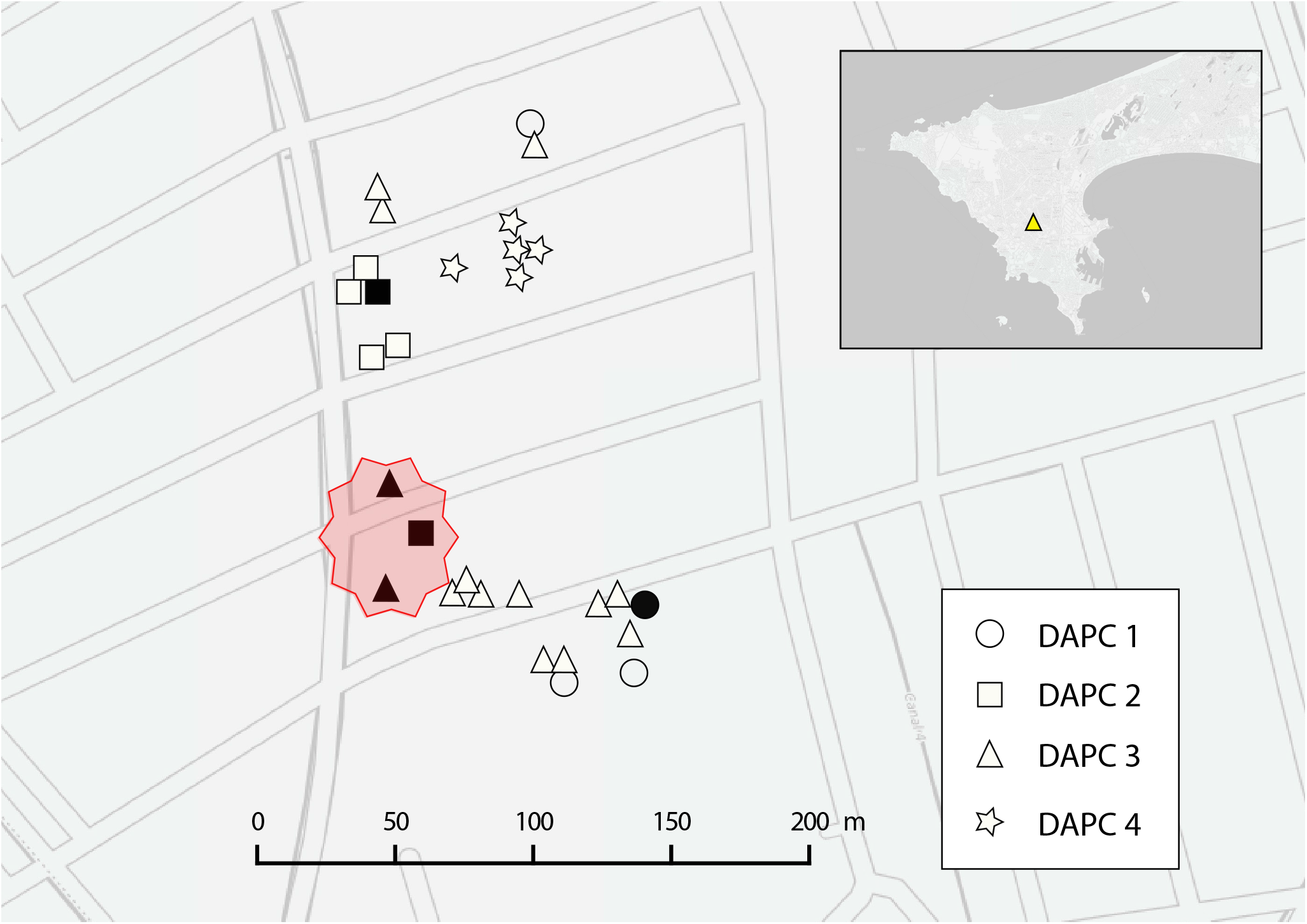
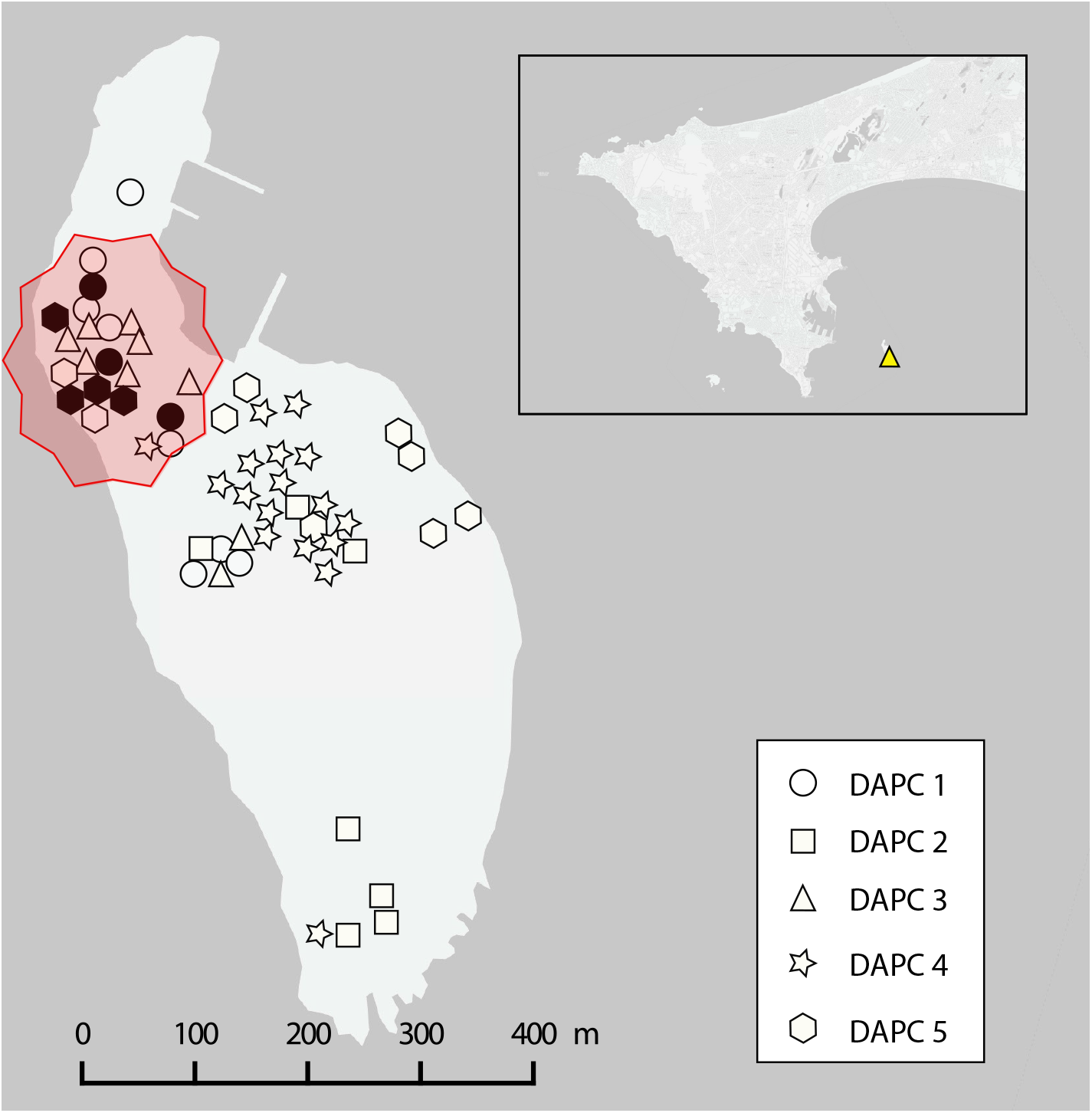
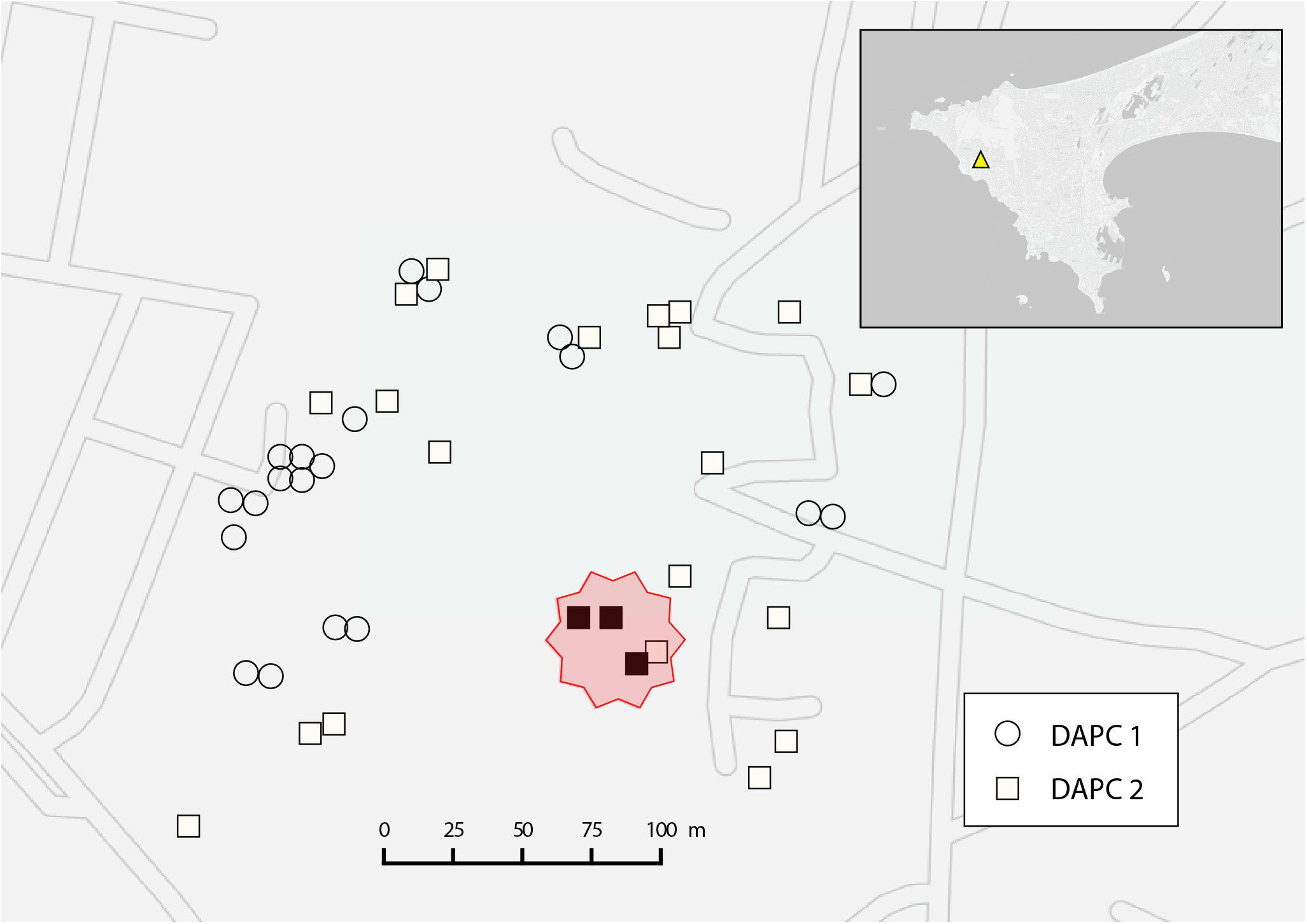
Sites in Dakar with possible over-prevalence zones of *Toxoplasma gondii* infection. Each small geometric shape represents the trapping localization of a given individual. Different geometric shapes were attributed to each DAPC groups. The infected individuals and the non-infected individuals are represented in black and white, respectively. The possible over-prevalence zones are surrounded by a circle with a dotted contour.

### 2.1. Spatial clustering analysis

Within each site, purely spatial cluster analysis was performed to test whether the infected mice were distributed randomly over space and, if not, to evaluate any identified spatial over-prevalence zone for statistical significance. Households (including shops and workshops) where rodent traps were set during the survey were considered as the smallest spatial unit in this analysis. ‘Spatial scan statistics’ was used to test the null hypothesis that the relative risk (RR) of *T. gondii* infection was the same between any household groups (or collection of household groups) and the remaining household groups of a sampling site. SaTScan software version 9.4.4 (Kulldorff, 1997, 2010), designed specifically to implement this test and using a Bernoulli model, imposed a circular scanning window on the map that moves across space. The area within the circular window, centered on the centroid of each household, varied in size from zero to a maximum radius never including more than 50% of the total population within a given site. The SaTScan software tested for possible over-prevalence zones within the variable window around the centroid of each household group. The number of Monte Carlo replications was set to 999, and over-prevalence zones with statistical significance of p <0.05 were reported.

### 2.2. Genetic structure analysis

Within each site where a spatial over-prevalence zone was identified, we investigated the possible occurrence of distinct genetic groups of mice using Discriminant Analysis on Principal Components (DAPC) (Jombart et al., 2010). DAPC was performed using the adegenet package (Jombart, 2008) for the R 3.4.0 software (R Development Core Team, 2009). This Bayesian clustering method is not based on a predefined population genetics model and is thus free from Hardy–Weinberg equilibrium assumptions.

## 3. Results

Purely spatial cluster analysis revealed three statistically significant possible over-prevalence zones in 3 sites of Dakar and on Goree Island.

A primary over-prevalence zone was identified in *Plateau-Reubeuss* (Figure 1-A). It contained all the 6 infected individuals within this site distributed over 5 households out of 28 total households in a 55 m radius (*P* = 0.014; SIR = 2.89 [95% CI: 1.06 – 6.31). In this site, 3 genetic groups of house mice were identified, (Figure 2-A), and 2 of them were found within the over-prevalence zone. From the 6 infected individuals within this over-prevalence zone, one individual belonged to the DAPC group 1 and 5 individuals belonged to the DAPC group 2.

**Figure 2.**
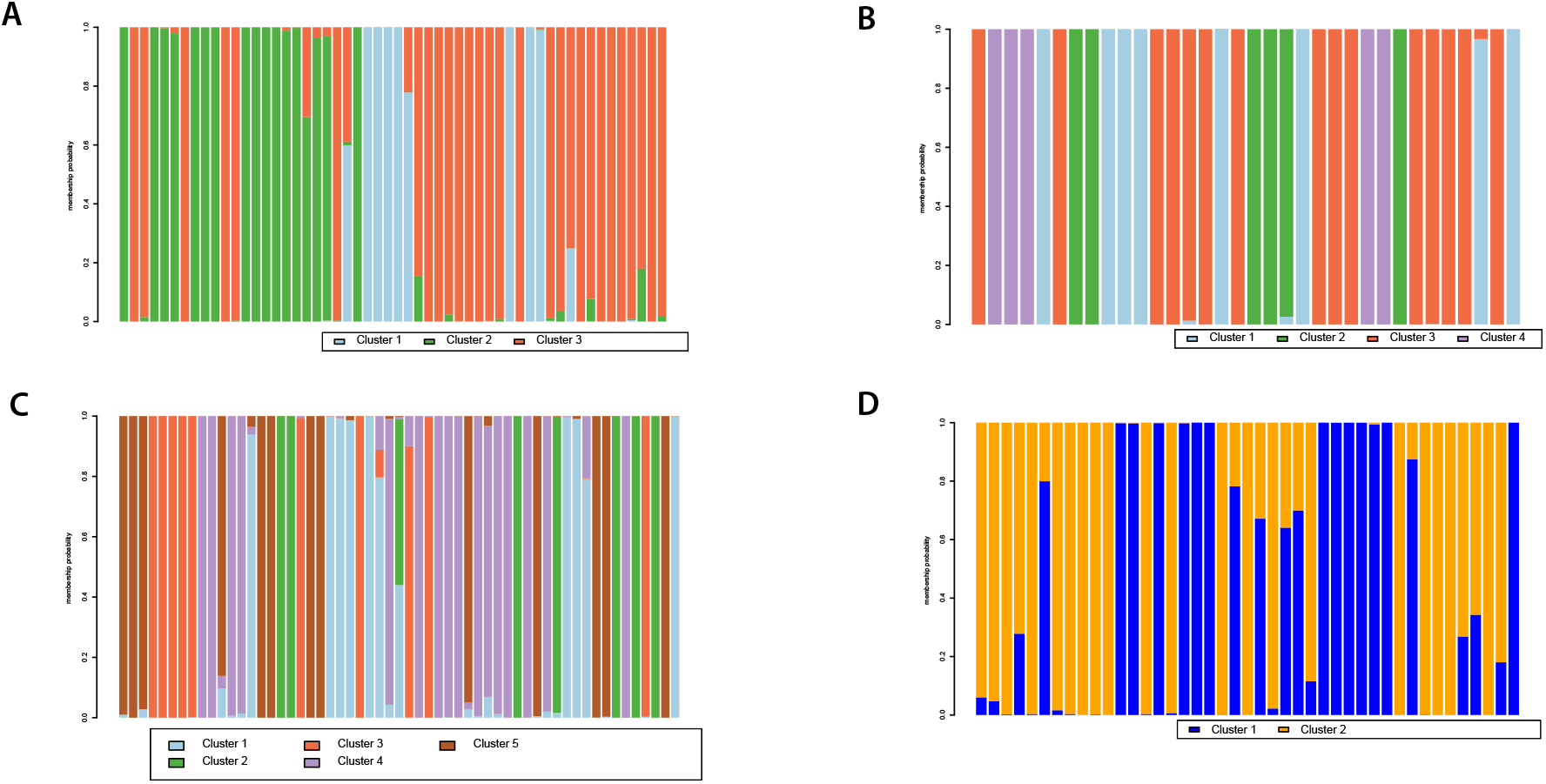
Clustering analysis with the Discriminant Analysis of Principal Components (DAPC) using the microsatellite genotypes of mice trapped in *Plateau-Reubeuss* (A), *Grand-Dakar* (B), Goree Island (C) and *Ouakam* (D). Individuals are aligned along the x-axis. Each barplot shows the probability on the y-axis (0.0–1.0) of an individual being assigned to a given number of clusters. Individuals are assigned either to one cluster (each cluster is represented by a different color) or to multiple clusters if their genotypes were admixed (indicated by multiple colors).

A second over-prevalence zone was found in *Grand-Dakar* (Figure 1-B). It contained 3 of the 5 infected individuals within this site distributed over 3 households out of 15 total households in a 40 m radius (*P* = 0.027; SIR = 5.77 [95% CI: 1.16 – 16.86]). In this latter site, 4 DAPC groups were identified (Figure 2-B), and 2 of them were found within the over-prevalence zone. From the 3 infected individuals within this over-prevalence zone, one individual belonged to the DAPC group 2 and 2 individuals belonged to the DAPC group 4.

A third over-prevalence zone was found on Goree Island (Figure 1-C). It contained all the 7 infected individuals identified on this island over 6 households out of 10 total households in a 97 m radius (*P* = 0.0069; SIR = 2.71 [95% CI: 1.09 – 5.59). Five DAPC groups were identified on this island (Figure 2-C), and 4 of them were found within the over-prevalence zone. From the 7 infected individuals within this over-prevalence zone, 3 individual belonged to the DAPC group 1 and 4 individuals belonged to the DAPC group 5.

The last over-prevalence zone was found in *Ouakam* (Figure 1-D). It contained all the 3 infected individuals identified in this site over 3 households out of 26 total households in a 18 m radius (*P* = 0.003; SIR = 10.71 [95% CI: 2.15 – 31.31). Two DAPC groups were identified in this site (Figure 2-D). The over-prevalence zone only included individuals from DAPC group 2.

## 4. Discussion

In the present study, we explored the occurrence of a possible association between the infectious status and the degree of kinship between individuals, which could give insights about the transmission route (oral or vertical) of *T. gondii.*

We investigated the occurrence of over-prevalence zones of *T. gondii* infection among mice from several sites in Dakar, Senegal. In several sites, the absence of detectable over-prevalence zones may be explained by gaps in sampling, the occurrence of more than one foci of oocyst contamination within the site or the movements of infected individuals within the site. In sites where an over-prevalence zone was identified, we investigated whether the infected mice in this over-prevalence zone belonged to a single genetic group — that we considered as probable family lines — or whether they belonged to more than one genetic group.

Our postulate is that finding an association between the degree of kinship and the occurrence of infection is not sufficient to conclude on the mode of transmission of *T. gondii* in mice. Indeed, given the limited proclivity of mice to move actively along large distances (Pocock et al., 2005), it is likely that mice of the same family would aggregate in space. Therefore, assuming a predominant role of oral transmission, the family line occurring on an area of recent oocyst shedding would be predominantly exposed to the oocysts found on the soil. A previous study investigating oocyst spatial distribution in an urban area has indeed demonstrated highly localized foci of oocysts’ occurrence, associated with cats’ defecation sites (Afonso et al., 2008). In particular, the sandy soil and dry climate of Senegal do not promote the spatial spread of oocysts or their long-term survival. In the case of vertical transmission, a spatial aggregation of the infected individuals could also be predicted as the transmission would occur only within a specific familial group that predominantly occupies a given area. In *Ouakam* for example, 3 infected mice gathered in the same over-prevalence zone and appeared to belong to the same DAPC group. Their infection could be due to an environmental contamination of this area by oocysts or to the occurrence of sibling mice infected through vertical route. However, finding individuals of distinct family lines in the same over-prevalence area and for which their infection status seems to be independent of their family line would be more suggestive of an oral route of infection.

The results provided in this study are mainly in accordance with the last scenario. Indeed, in 3 of 4 sites where possible over-prevalence zones were identified (*Plateau-Reubeuss*, *Grand-Dakar*, and Goree Island), the infected individuals within those over-prevalence zones were distributed over more than one DAPC group, supporting the hypothesis that infection occurrence is more related to spatial location than to family effects. These results suggest that the acquisition of infection in mice from Dakar is independent of kinship and is hence more likely to occur through an oral route.

This conclusion appears to be in contradiction with a number of field studies in which the results supported a vertical transmission of *T. gondii* in several species of rodents (Murphy et al., 2008; Thomasson et al., 2011; Webster, 1994). In those previous studies, high prevalence levels of *T. gondii* infection were reported among rodents in areas that appear to be relatively free of cats. These findings provide indirect evidence that vertical transmission can be sufficient to the perpetuation of transmission. In line with this, a field study was performed on wood mice *Apodemus sylvaticus* in a location where low cat density is reported (Bajnok et al., 2015). The wood mice from this location were found to belong to four distinct family lines which were distributed in different parts of the study zone. The prevalence of infection was found to be significantly different in each of the families and was linked to host genotype rather than location of capture. This suggested that parasite infection was indeed associated with families and non-randomly distributed throughout the families, which is in accordance with a predominantly vertical route of transmission.

However, all these studies were conducted in the United Kingdom, limiting the diversity of epidemiological situations in the exploration of this issue. Indeed, in situations where cats densely populate an area — as it is the case in Dakar, Senegal (Bend, 1980; Lahamdi, 1992) — and in which rodents are therefore heavily exposed to oocysts, the high predation pressure would reduce the rodents’ overall lifespan and the likelihood of reaching sexual maturity to transmit the infection to offspring through vertical transmission (Turner et al., 2013). In particular, assuming a role of parasitic manipulation in facilitating the predation of rodents by cats, a primary involvement of vertical transmission could be detrimental for the sustainability of the cycle as infected individuals would be less likely to reach sexual maturity and reproduce compared to non-infected individuals as they would die earlier in age. Finally, the geographical variability of the *T. gondii* strains can also have an important role on the transmission dynamics of *T. gondii* (Galal et al., 2018a). Virulence variability of *T. gondii* strains could indeed impact the rate of vertical transmission as suggested by a predictive model by (Lélu et al., 2013).

Altogether, the factors quoted above emphasize on the importance of considering the specificities of each epidemiological situation in inferring on transmission dynamics from empirical data. Additional research is needed to further explore this issue in different epidemiological situations and for different rodent species.

## Acknowledgements

We thank Karine Berthier for helpful discussions. We thank the French Agence Nationale de la Recherche (ANR project IntroTox 17-CE35-0004), the University of Limoges and the Nouvelle-Aquitaine region for funding this research.

